# Development of a Platform for High-Resolution Ion Mobility Separations Coupled with Messenger Tagging Infrared Spectroscopy for High-Precision Structural Characterizations

**DOI:** 10.1101/2024.09.06.611750

**Authors:** Christopher P. Harrilal, Sandilya V.B. Garimella, Randolph V. Norheim, Yehia M. Ibrahim

## Abstract

The ability to uniquely identify a compound requires highly precise and orthogonal measurements. Here we describe a newly developed analytical platform that uniquely integrates high resolution ion mobility and cryogenic vibrational ion spectroscopy for high-precision structural characterizations. This platform allows for the temporal separation of isomeric/isobaric ions and provides a highly sensitive description of the ion’s adopted geometry in the gas phase. The combination of these orthogonal structural measurements yields precise descriptors that can be used to resolve between and confidently identify highly similar ions. The unique benefit of our instrument, which integrates a structures for lossless ion manipulations ion mobility (SLIM IM) device with messenger tagging infrared spectroscopy, include increased resolution and the ability to record the IR spectra of all ions simultaneously. The SLIM IM device, with its 13m separation path length, allows for multipass experiments to be performed for increased resolution as needed. It is integrated with an Agilent qTOF MS where the collision cell was retrofitted with a cryogenically (30 K) held TW SLIM device. The cryo-SLIM is operated in a novel manner that allows ions to be streamed through the device and collisionally cooled to a temperature where they can form non-covalently bound N_2_ complexes that are maintained as they exit the device and are detected by the TOF mass analyzer. The instrument can be operated in two modes: IMS+IR where the IR spectra for mobility-selected ions can be recorded and IR-only mode where the IR spectra for all mass-resolved ions can be recorded. In IR-only mode, IR spectra (400 cm^-1^ spectral range) can be recorded in as short as 2 seconds for high throughput measurements. This work details the construction of the instrument, modes of operation, and provides initial benchmarking of CCS and IR measurements to demonstrate the utility of this instrument for targeted and untargeted approaches.

## INTRODUCTION

Mass spectrometry (MS) based measurements have become a mainstay for the identification and quantification of the contents of complex mixtures, especially in -omics-related fields.^1-5^ MS excels in measurement speed, accuracy, sensitivity, and precision, allowing molecules to be characterized quickly compared to other non-MS-based methods. Analyte identification generally consists of measuring the properties of an unknown analyte and comparing them to the measured properties of known compounds. This approach is known as library matching and relies on the large-scale characterization of standard compounds.^6^ Libraries of MS-based techniques can contain measurements of mass, fragmentation spectra (i.e., MS^n^), collisional cross sections (CCS), as well as other metrics such as retention time when using gas or liquid chromatography (GC/LC).^6-9^

The ability to uniquely characterize a molecule relies on precise orthogonal measurements.^9, 10^ Mass spectrometric measurements provide a level of precision that can differentiate compounds based on the number and type of atoms they contain within the mass error of the measurement. High-resolution mass measurements often have uncertainties of a few parts per million (ppm).^11^ Differentiating the chemical space within a given mass tolerance relies on other orthogonal measurements such as MS^n^, CCS, or GC/LC retention times.^9, 12, 13^ These measurements probe how a molecule dissociates, the overall size of the molecule, and the retention time through an LC (or GC) column. In general, the greater the number of orthogonal measurements matched between an unknown analyte and a library entry, the more confident the molecular annotation.^14^ Areas that remain challenging in molecular characterization are the distinction between isomeric or isobaric compounds that share similar fragmentation patterns or the identification of small molecules that fail to provide diagnostic fragments using conventional activation techniques. In some scenarios, LC/GC may aid in distinguishing isomers. Still, these separation techniques do not have enough resolving powers, are not broadly applicable, or require tailored method developments.^15-17^

A more generalizable approach to resolving between compounds of this type is to leverage measurements sensitive to an ion’s 3-D shape. These measurements are useful as the 3-D shape adopted by an ion in the gas phase strongly depends on its arrangement of atoms.^18^ Ion mobility measurements serve as one type of probe of a molecule’s 3-D shape. These measurements are made by monitoring an ion’s drift time through a buffer gas cell under the influence of a weak static or oscillating electric field.^19, 20^ The drift time is proportional to the ion’s mobility through equation 1:

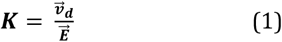

where 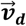 is the net ion relative velocity through the buffer gas in a steady-state condition and 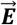 is the electric field. The mobility is often expressed as the momentum transfer cross section (Ω) through the Mason-Schamp equation^21^ (eq.2):

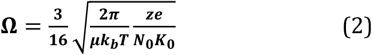

where µ is the reduced mass of the ion and buffer gas pair, ***k***_***b***_ is the Boltzmann constant, T is the buffer gas temperature, ***z*** is the ion charge, ***e*** is the elementary charge, ***N***_**0**_ is Loschmidt’s number, and ***K***_**0**_ is the reduced mobility 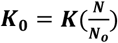, used to normalize for experimental conditions, where ***N*** is the gas number density at experimental conditions. Ω is a term that quantifies the average effective area that interchanges momentum per unit time with the buffer when there is a velocity difference between the ion and buffer gas^22^ and is often used as a numerical description to compare relative ion sizes.

Current commercially available ion mobility spectrometry (IMS) platforms achieve resolving powers (R_p_) of 300-400^23, 24^ and can differentiate between ions that differ in Ω by 0.5 % or less.^25^ With this level of resolving power isomeric glycans, lipids, and various metabolites can be resolved, aiding in compound identification in untargeted approaches.^26-30^ Much higher levels of separation power are achievable with traveling wave ion mobility spectrometry (TWIMS) based systems where ions can be rerouted through the device for extended separations.^31-35^. Under these conditions, ions with highly similar 3-D structures can be resolved or partially resolved, making IMS-based methods more attractive for complex samples. In extreme cases isotopomers, molecules with identical 3-D geometries but differing arrangement of nuclei, have been separated by mobility^36-38^ in a predictable manner.^39-41^ The resolving powers in these experiments ranged from ∼1500-2000, demonstrating the level of separation that can be achieved and the future of IMS for high-precision measurements.^30^

Additional probing of a molecule’s 3-D shape in the gas phase can be performed using action spectroscopy.^42^ In these approaches, light initiates a mass change of isolated ions, allowing absorption bands to be probed. One technique to record IR spectra is infrared multiple photon dissociation (IRMPD)^43^, as demonstrated by Free Electron Laser for Infrared eXperiments (FELIX)^44-46^ and Infrared Laser Center of Orsay (CLIO)^47^ institutions. With this technique, an ion is mass isolated, stored in an ion trap, and irradiated with a narrow bandwidth and high-power IR light of a certain frequency. When the frequency of IR light is in resonance with a vibrational transition, the ion absorbs multiple photons until its internal energy is sufficiently high to dissociate on a millisecond time scale.^45^ The abundance of fragment ions is detected as a function of the frequency of the IR light as it is scanned through a spectra range, allowing the IR spectrum of the ion to be recorded. This technique excels in probing smaller, non-flexible molecules, however, this approach is prone to a phenomenon known as vibrational transparency, which affects flexible and non-covalently bound systems, where vibrational bands can appear missing or with skewed intensities from the recorded spectrum.^48-50^ Recent work has incorporated IRMPD into analytical workflows that include LC/MS and MS^2^ measurements for de-novo structural elucidation.^51, 52^

An alternative technique to record IR spectra is messenger tagging.^53, 54^ In this method, mass (or mobility) -isolated molecules are trapped in an ion trap and cooled to temperatures of ∼30-50^°^K.^55, 56^ At these temperatures, N_2_ can condense on the ion forming a non-covalent complex causing a mass increase of the isolated ion by 28 Dalton. By irradiating the cryocooled ion packet with IR light in resonance with a vibrational transition, the ion will gain enough internal energy to break the weak non-covalent interaction with N_2,_ resulting in a detectable mass change. With this approach a limited number of photon absorption event is enough to create the detectable “action” and minimizes spectral distortion associated with a multiphoton process. Further, the cryogenic temperatures limit thermal broadening, yielding well-resolved vibrational bands.

The IR spectra serves as a structural fingerprint of the ion as it is a direct report of the hydrogen bonding network^57, 58^ and provides complementary information to an IMS measurement. In many cases the IR spectrum recorded using messenger tagging can serve as a precise descriptor making it useful for compound identification and annotation but has not been strongly considered as an analytical technique due to the low throughput and technical requirements. Pioneering work by Rizzo and co-workers, however, has pushed the messenger tagging technique as a viable analytical technique. The Rizzo group has developed analytical platforms that couple either TWIMS using Structures for Lossless Ion Manipulations (SLIM) (allowing for collision induced dissociation and IMS^n^)^59, 60^ and/or LC-MS with messenger tagging demonstrating that IR spectra can be recorded in a matter of seconds.^56, 61^ The increase in speed compared to more traditional approaches is largely due to their ability record IR spectra for multiple isomers simultaneously and the use of a continuous wave (CW) laser.^56, 62, 63^ The utility of their approach for isomer identification has been demonstrated by resolving between glycan isomers and several other metabolites.^64, 65^ In their approach SLIM separations provide temporal separation of isomeric species allowing for highly precise IR signatures to be recorded for each compound and stored for future database matching.^59^

In the following work we further the development of high-resolution IMS and messenger tagging as an analytical technique and detail the development of a platform that couples high-resolution TWIMS using SLIM and messenger tagging for highly precise structural characterizations of gas phase analyte ions. This instrument is designed for detailed complementary structural characterizations that can be used for molecular identification and de-novo structural elucidation when used in conjunction with high-accuracy mass and MS^n^ measurements and corresponding predictive tools. This manuscript focuses especially on describing the experimental platform, advancements in methodologies, and demonstrating the combined powers of high-resolution CCS and cryogenic IR measurements.

## METHODS

### Instrument Overview

The experiments were performed on a modified Agilent qTOF-MS (model 6538, Agilent Technologies, Santa Clara, CA), as shown in Figure 1. Ions are generated through nano-electrospray ionization and guided into a stainless-steel heated capillary held at 130°C. Desolvated ions enter a dual funnel interface. The high-pressure ion funnel was held at ∼ 8 torr while the second funnel was held at 2.5 Torr. Ions were collimated and introduced into a 13-meter TW-SLIM device, held at ∼ 2.5 Torr of pure N_2_. Ions can be continuously introduced to the device, where no mobility separation occurs, or pulsed for IMS experiments. For IMS experiments, a pulse of ions is introduced by modulating the DC bias of a single funnel electrode to control the number of ions injected into SLIM. Ions are accumulated in the first section of SLIM using a TW of low amplitudes (square profile, 5 V_pp_). Following ion accumulation, the TW amplitude is then increased to the separation TW (32 V_pp_). The details of this SLIM device have been described previously.^66^ Briefly, ions can be routed once through the device, resulting in a 13-meter separation, or cycled for a user-defined number of cycles for extended separations. Ions exit the SLIM device into a rear-ion funnel where they are collimated and introduced to a constant-field (CF) SLIM module held at ∼300 mTorr. The CF-SLIM was designed to turn the ion beam by 90° (see Supplementary Information Figure S1) as it exits the TW-SLIM and enters the Agilent 6538 system. The standard Agilent ion source housing was removed and replaced with the vacuum housing of the CF SLIM. The 90^°^ CF SLIM interface grants optical access for the laser via a 0.5-inch diameter CaF_2_ window mounted onto the vacuum housing. The window is centered vertically and horizontally between the two CF-SLIM boards allowing the laser (continuous wave 3300-3900 cm^-1^, CLT series from IPG Photonics, USA) to have a clear line of sight through the qTOF MS. The design of the CF-SLIM has been discussed previously.^67, 68^ Briefly, the adjacent rung electrodes are supplied with alternating phases of RF and are resistively coupled to allow for a DC gradient to be formed between the entrance and exit of the device. Resistively coupled DC-only guard electrodes sit on both sides of the rung electrodes, and are 2 mm long, spanning 2 rung electrodes. The guard electrodes prevent ions from escaping laterally from the device. These electrodes are held at ∼5-10 V above the rung electrode potential. An electric field of 3.34 V/cm moves ions through the CF SLIM. After the CF-SLIM, ions enter the Agilent 6538, passing through a series of lenses, including a skimmer, octupole, and selection quadrupole, before entering the cryogenically cooled TW-SLIM. The TW-SLIM trap replaces the original collision cell in this instrument, Supplementary Information Figure S2 and Figure S3 show the schematic diagram and some of the dimensions for one of the PCBs and a cross-section of the assembly, respectively. The TW-SLIM is mounted on the existing mounts for the original collision cell using parts designed from PEEK material. The PCB for this SLIM is made from Rogers RO4350B material (woven glass reinforced hydrocarbon/ceramics) and measured 179 mm long, 60 mm wide, and 1.6 mm thick. The electrical connections between electrodes within a single PCB are moved to an inside layer buried inside the board such that a smooth surface exists on the non-electrode side of each board. This allows for a smooth surface in contact with the bottom and top oxygen-free copper mounting blocks, allowing for efficient heat transfer. The top and bottom boards are electrically connected via a pogo-type Molex connector that compresses between the boards when mounted within the copper housing. The boards are spaced by 3 mm copper spacers, which contain 4 alignment pins to center the boards in the mount. Figure S2 shows the board layout, which contains 6 RF electrodes and 5 segmented traveling wave electrodes and flanked by two guard electrodes. Cooling the cryo-TW-SLIM is accomplished using a single-stage He cryostat (CH-110; Sumitomo, JPN) with an ultimate temperature of 20 K. The contact between the SLIM and He cryostat is accomplished by a copper finger directly mounted to the He cryostat and contacts a receiving copper piece directly mounted to the top of the SLIM copper block housing. The TW-SLIM is held at ∼ 30 K for messenger tagging experiments. The temperature of the Cryo-TW-SLIM can be controlled above 20 K using two heater cartridges (Hotwatt 11 Ω wired in series, each rated for 24 W power output when 24 V is applied), a temperature sensor (Silicon diode, part number DT-670C-CU, LakeShore, Westerville, OH), and a temperature controller (model 335, LakeShore. Wester-ville, OH). The top lid of the Agilent vacuum manifold is modified so that an extension chamber can be mounted on the lid to support the He cryostat. Ultra-high purity He and N_2_ are mixed and introduced into the SLIM device via a 0.25 in i.d. stainless tube suspended directly above the gas port of cryo-TW-SLIM module (Supplementary Information Figure S3). A hole in the copper block and top TW-SLIM board allows gas to enter the device. The copper blocks and spacers encapsulate the SLIM device, reducing the rate at which the gas can exit the device. Ions enter and exit the cryo-TW-SLIM through circular orifices of i.d. 3 mm. The cryo-TW-SLIM replaces the Agilent collision cell with no additional lenses before or after the cryo-TW-SLIM module. The MIPS power supply provided all cryo-TW-SLIM’s DC, RF, and TW voltages (GAA Custom Electronics LLC, Kennewick, WA). Ion spectroscopy experiments used a mixture of He/N_2_ estimated to be in a 90/10 ratio, while non-ion spectroscopy experiments used only He. A final operating pressure of ∼3×10^−5^ Torr (measured outside the SLIM trap) is achieved to stream ions through the device while the base pressure was ∼10^−7^ Torr when no gas was used in the cryo-TW-SLIM. Ions exit the cryogenically held TW-SLIM and pass through two transport hexapoles into the time-of-flight (TOF) flight tube for detection. The signal from the Agilent MCP was routed to a computer hosting a U1084, 8-bit analog-to-digital convertor (Acqiris, Plan-les-Ouates, Switzerland) and processed through a control software developed in-house.

**Figure 1.**
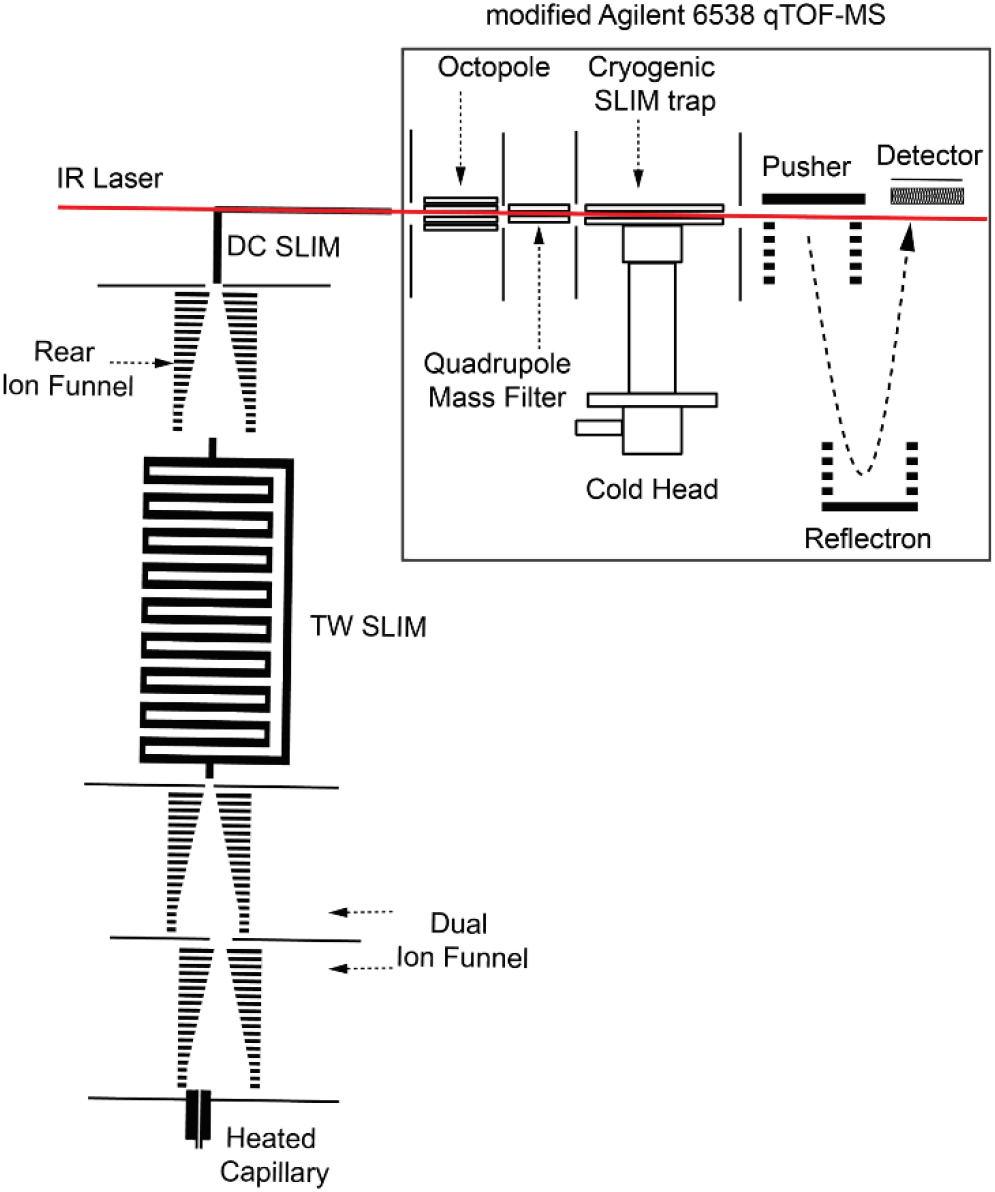
(a)Schematic of SLIM-Cryogenic IR instrument retrofitted from an Agilent 6538 qTOF MS.

### Messenger Tagging

To record infrared spectra, we use the messenger tagging technique. Generally, ions are introduced into an ion trap held at cryogenic temperature. Before ions introduction, a pulse of a gas mixture (He/N_2_) is introduced to the ion trap. As ions enter, they experience collisions with the gas and thermalize to the trap temperature. Some N_2_ molecules form non-covalent bonds with the ions as the gas is pumped away, forming N_2_ ‘tagged’ ions. Once the gas is pumped away, these ions are gently extracted from the trap and detected. A portion of the ion population remains untagged and is detected as an [M+nH]^n+^ ion while the tagged population is detected as an [M+nH+m28]^n+^ where ‘m’ represents the number of N_2_ tags adducted to the ion and ‘n’ represents the number of protons. The ratio of tagged to untagged ions can be modulated by irradiating the ion packet before it is ejected from the ion trap. If the laser wavelength is resonant with a vibrational transition of the cryocooled ion, the ion will absorb a photon and gain enough internal energy to break the non-covalent interaction between the N_2_ and molecule. By plotting the ratio of [M+tag]/([M]+[M+tag]) as a function of the laser wavelength it is possible to obtain the IR spectrum of the selected ion.

In our instrument, we follow a slightly different approach. Rather than trapping ions, we gently stream ions (i.e., no ion trapping) through the cryogenically held (30 K) TW-SLIM. As the precursor ions transverse the TW-SLIM device, they undergo several collisions and retain one or more tags as they exit the device and enter the TOF pusher region. Low-flow needle valves (part number SS-SS4-KZ, Swagelok) are used to independently regulate the flow of N_2_ and He. The composition of the mixture is roughly 90/10 He/N_2_ but can be adjusted to obtain an ideal tag-to-parent ratio. Operation in this manner makes the experiment more streamlined and increases the ease of operation as timings associated with ion trapping and gas pulsing are non-factors. Further operation in this manner allows for all mobility-separated ions to be probed simultaneously. Two modes of recording IR spectra are used in this work: IMS+IR and IR-only mode. The IMS+IR is leveraged to measure the IR spectra of mobility-resolved isomers. The IR-only mode allows a higher throughput (∼2-30 seconds) spectroscopic probe of mass-resolved ions.

### Data Processing

MS and IMS data are recorded in the universal ion mobility format (.UIMF) and viewed in the PNNL UIMF Viewer (https://github.com/PNNL-Comp-Mass-Spec/UIM-FViewer) file format where each frame represents a single instrument acquisition cycle and is processed using PNNL-developed preprocessor tools.^69^ The laser is synchronized to the start of the experiment and the wavelength is recorded at the beginning of each frame. Raw data are aligned using a linear drift time aligner (https://github.com/PNNL-Comp-Mass-Spec/IMS-Drift-Time-Aligner) to account for pressure-related fluctuations.^70^ Intensity values for a precursor ion and the associated tagged mass of interest are pulled from a user-defined mobility window from the UIMF file as a function of the frame. Intensity values are plotted as a ratio of [M+H+tag]^+^ / ([M+H]^+^+[M+H+tag]^+^) for each frame. The frame number is then converted to the laser wavelength to plot the final IR spectrum.

### Sample

All samples were purchased from Sigma Aldrich. A mixture of YGGFL, SDGRG, and GRGDS was prepared with a final concentration of 10 µM for YGGFL and 20 µM for the reverse peptides. Samples were diluted using 50/50 ACN/H_2_O.

## Discussion

The evaluation and use of this TW-SLIM platform for mobility separations have been published elsewhere.^66^ Two modes of operation are available to record IR spectra: IMS+IR mode and IR-only mode. In IMS+IR mode (the time sequence is shown in Supplementary Information Figure S4), ions are first streamed into the SLIM device for a user-defined fill time, typically 100-500 ms. During this time, the TW amplitude is set between 2-5 volts to accumulate ions. Once the fill time has ended, the TW amplitude is increased to the separation amplitude and ions begin to mobility-separate. At the end of the 13 m path length, ions can either be directed out of the SLIM or cycled a defined number of times. Ions exiting SLIM are directed to the cryogenic TW-SLIM. During a tagging experiment, the TW conditions in the cryogenic SLIM determine whether the precursor ions retain any N_2_ tags as they pass through the device. This is demonstrated in Figure 2 which shows the arrival time distributions (ATDs) and corresponding mass spectra of [YGGFL+H]^+^, [GRGDS+H]^+^, and [SDGRG+H]^+^ using a 100 ms in-SLIM accumulation (TW = 5 V_pp_) followed by a separation condition of 32 V_pp_ and speed of 208 m/s, but with varied TW amplitudes in the cryo-SLIM module. Figure 2a shows the corresponding arrival time distribution when the TW amplitude of the cryo-SLIM is set to 2 V_pp_. The ATD is as expected with typical peak widths (FWHM) of ∼2.5 ms, providing enough resolving power to near baseline separate the protonated reverse peptides GRGDS and SDGRG, which have collisional cross-section differences of ∼1.5%. CCS values for [SDGRG+H]^+^, [GRGDS+H]^+^, and [YGGFL+H]^+^ are measured as 202.1, 205.3, and 227.24 Å^2^, respectively, and are obtained through calibration using Agilent tuning mix ions (see Supporting Information for the calibration procedure). The measured CCS values are within 1% of reported CCS values.^7, 23^ The corresponding MS for this separation is shown in Figure 2b, which shows no tagged precursor ions. The results shown in Figure 2a indicate that a TW amplitude of 2 Vpp produces an effective temperature sufficient to dissociate the N_2_-ion complex. Figures 2c and 2d show the corresponding ATD and mass spectrum of the peptide mixture when the TW amplitude of the cryo-SLIM is set to 0 V_pp_. Figure 2d shows both precursor masses retain 2 or more tags as they pass through the cryo-SLIM. The reverse peptide precursor mass picks up two tags, with the first tag in a near 1:1 ratio and the second tag in a 1:0.5 ratio with the precursor mass. On the other hand, [YGGFL+H]^+^ picks up 3 tags, with very little precursor ion remaining. The first and second tags are nearly equal in abundance and about 12× the abundance of the precursor, while the third tag is about half the abundance of the first 2 tags. Under these conditions, the IR spectrum can be taken for [YGGFL+H]^+^ as this ion is resolved by mass and mobility from all other ions in this mixture. However, the mobility separation between [GRGDS+H]^+^ and [SDGRG+H]^+^ is not retained as the pair transverses the cryo-SLIM (Figure 2c). This is due to the lack of an axial electric field to move ions inside the cryo-TW-SLIM at 0 V_pp_. Under these conditions the ions rely on their incoming translational kinetic energy to transverse the device. On the other hand, the absence of the TW electric field allows ions to become thermalized to the buffer gas temperature, tagged, and gently transferred out of the cryo-SLIM. This process makes it possible to record their IR spectra. However, the lack of the electric field also results in the broadening of the ATD for each peak. The FWHM of the YGGFL peak increases from 2.47 ms (Figure 2a) to 5.48 ms (Figure 2c). A similar level of peak broadening occurs for the reverse peptides, causing them to become non-resolved under these conditions. To remedy this problem, the reverse peptide pair can be rerouted in the 13-m SLIM to further increase their separation in time before being introduced to the cryo-SLIM. This is shown in Figure 2e, where the reverse peptide ions are re-routed for an additional pass prior to exiting the 13-m SLIM. The ATD for [YGGFL+H]^+^ remains unchanged as this ion exited the SLIM without undergoing an additional pass. As a benchmark, the ATD in Figure 2e is recorded first with the TW amplitude in the cryo-SLIM set to 2 V_pp_, allowing this separation to be directly comparable to that shown in Figure 2a. In comparing the ATDs of the reverse peptide pair between Figure 2a and Figure 2e, we see that the temporal separation between the reverse peptides increases from 6.83 ms to 13.36 ms. For completeness, the mass spectrum associated with this separation is shown in Figure 2f, which, as expected, does not show any tag retention for the precursor ions. Figure 2g shows the resulting ATD of [YGGFL+H]^+^ and the reverse peptide pair when the TW amplitude in the cryo-SLIM is set to 0 V_pp_ and the reverse peptide pair is re-routed for an additional pass. Again, when the TW amplitude is set to 0 Vpp, the ATDs broaden. However, because the reverse peptides are further separated, they are still resolved in time. The corresponding mass spectrum of this separation is shown in Figure 2h, which is similar to that in Figure 2d, where multiple tags appear in the absence of the TW electric field. Under these conditions, the IR spectrum of the reverse peptides can be taken as they are resolved in mass and mobility from all other ions in the mixture.

**Figure 2.**
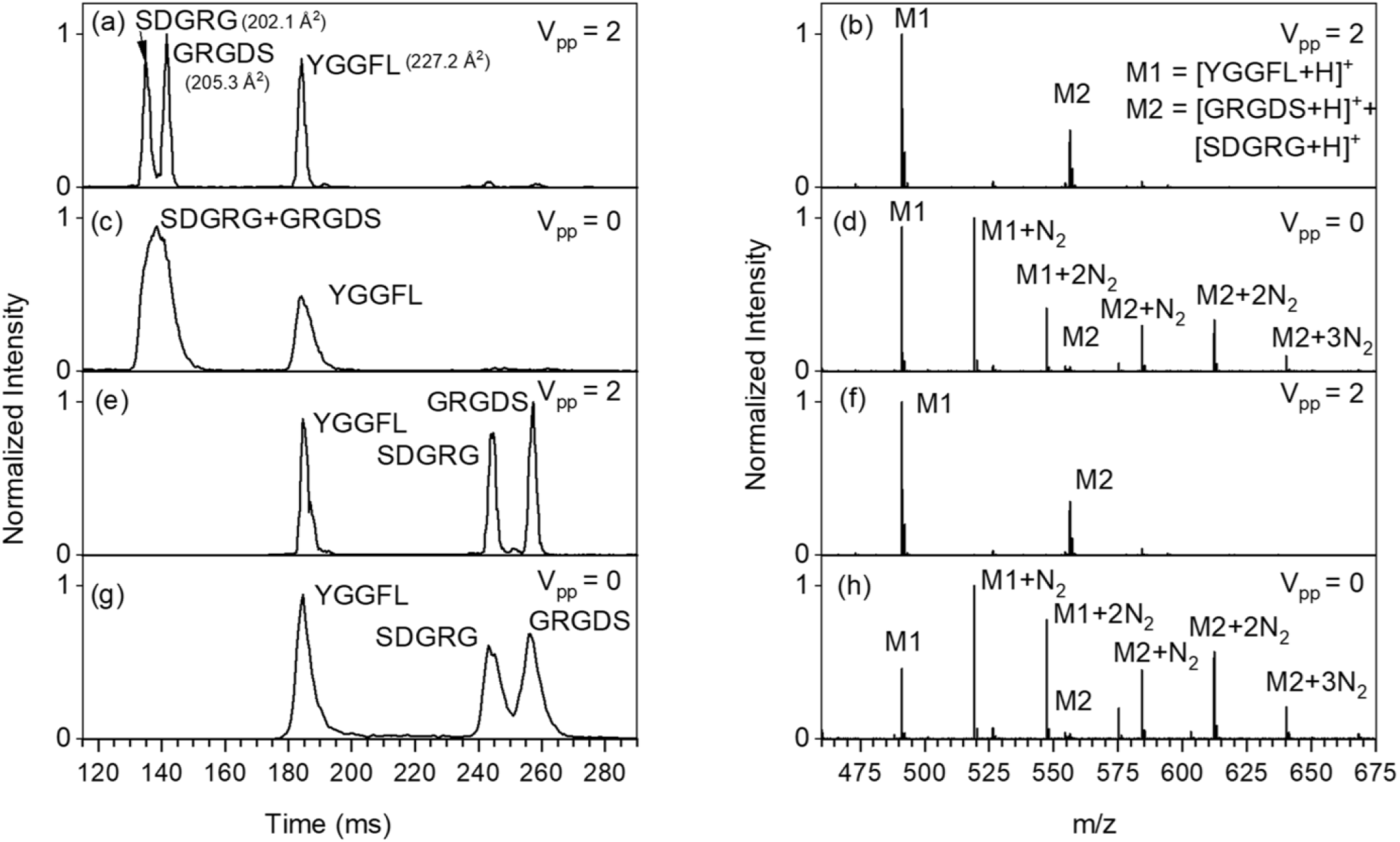
Arrival time distributions of the peptide ions [YGGFL+H]^+^, [GRGDS+H]^+^, and [SDGRG+H]^+^ (a, c, e and f) and their associated mass spectra (b, d, f, and h). All experiments used a 100 ms fill time (TW amplitude = 5 V_pp_) followed by a 32 V_pp_ TW amplitude for separation within the 13-m SLIM. The arrival time distributions shown in (e) and (g) correspond to performing an additional cycle within the 13-m SLIM for the reverse peptide pair prior to routing to the cryo-SLIM. A guard voltage of 6 V was used in the Cryo-TW-SLIM.

To understand the effect of the cryo-SLIM’s guard electrode on the preservation of the tagged species, their intensities were recorded as a function of the guard voltage and the corresponding precursor’s intensity (Supplementary Information Figure S5). This was done for the precursor *m*/*z* of 556 and 491. These results indicate a guard of 6 V provides the best ratio of tagged to precursor ions. At guard values below 6 V, fewer ions pass through the cryo-SLIM due to insufficient ion confinement. At guard values above 6 V, fewer tagged ions remain due to ions being forced towards the two SLIM surfaces causing them to experience higher RF oscillations and higher internal energy.

Figure 3a shows the data when recording the IR spectrum of all mobility-separated ions in the sample using a laser scan rate of ∼1 cm^-1^/acquisition and acquisition length of 820 ms and using 500 ms fill time. Figure 3a shows the arrival times of all ions on the x-axis and their *m*/*z* values on the y-axis. The intensity is shown as a color scale. The IR spectra are recorded using the same TW conditions as in Figure 2g, a TW amplitude of 32 V_pp_ for the 13-m SLIM separation, re-routing of the reverse peptide pair for an additional pass, and a TW amplitude of 0 V_pp_ in the cryo-SLIM. The only difference is the IR spectra for Figure 3 were recorded using a 500 ms fill time as opposed to a 100 ms fill time used in Figure 2. Figure 3a show 4 prominent ions, [YGGFL+H]^+^, [GRGDS+H]^+^, [SDGRG+H]^+^, and [SGRG+H]^+^. The [SGRG+H]^+^ ion (*m*/*z* = 404.19) is a y_4_^+^ fragment of [GRGDS+H]^+^ most likely formed within the ion source, as it is mobility-separated from its precursor ion. Each precursor ion labeled is resolved by mobility or mass from all other species present and forms 2 or more tags as it passes through the cryo-SLIM. Under these conditions, the IR spectrum of all species was simultaneously recorded by monitoring the intensity of the tagged and precursor ions in a defined mass and mobility window. The intensity of all tags is considered when plotting the IR spectra. The IR spectra for the 4 prominent ions are shown in Figure 3b-e. The accuracy of our IR spectra can be benchmarked by comparing the IR spectrum of [YGGFL+H]^+^ to work previously published.^71, 72^ Further, the modest effect on the resulting IR spectrum when considering multiple tags vs. a single tag is demonstrated in Supplementary Information Figure S6. The IR spectra of [GRGDS+H]^+^ (Figure 3c) and [SDGRG+H]^+^ (Figure 3d) appear quite different from one another and both contain several vibrational bands. The complexity of these spectra can be largely attributed to the presence of arginine, which contains 5 vibrational transitions that may appear in this wavelength region in addition to the 4 backbone amide NH stretches and 2 OH functional groups. The complexity of these 5 amino acid containing peptides can be contrasted to [YGGFL+H]^+^, which only contains 4 backbone amide NH stretches and 1 OH stretch. Further, the IR spectrum of the y_4_^+^ fragment of [SDGRG+H]^+^ is also recorded (Figure 3e), highlighting the ease of recording IR spectra when tagging occurs without pulsing gas and trapping ions. In this approach, the IR spectrum of any ion in the sample can be recorded with little experimental effort. The IR spectrum of [DGRG+H]^+^ appears quite complex, with several transitions appearing in the 100 cm^-1^ span between 3500 and 3600 cm^-1^. This spectrum’s complexity may indicate several similar structures that are not fully resolved by mobility. Additional mobility passes were not performed for this fragment ion. The IR spectra shown in Figure 3 were recorded using a step size of about 1 cm^-1^ per acquisition cycle for a total acquisition time of ∼5 minutes. Larger step sizes and shorter fill times can be used to acquire IR spectra in much shorter times. Supplementary Information Figure S7 shows the IR spectrum of [YGGFL+H]^+^ as a function of the laser step size from 1-7 cm^-1^. Using a step size of ∼3 cm^-1^ / acquisition cycle, the IR spectrum can be reasonably reproduced with a total acquisition time of ∼2 minutes but at the cost of slightly less resolved spectra.

**Figure 3.**
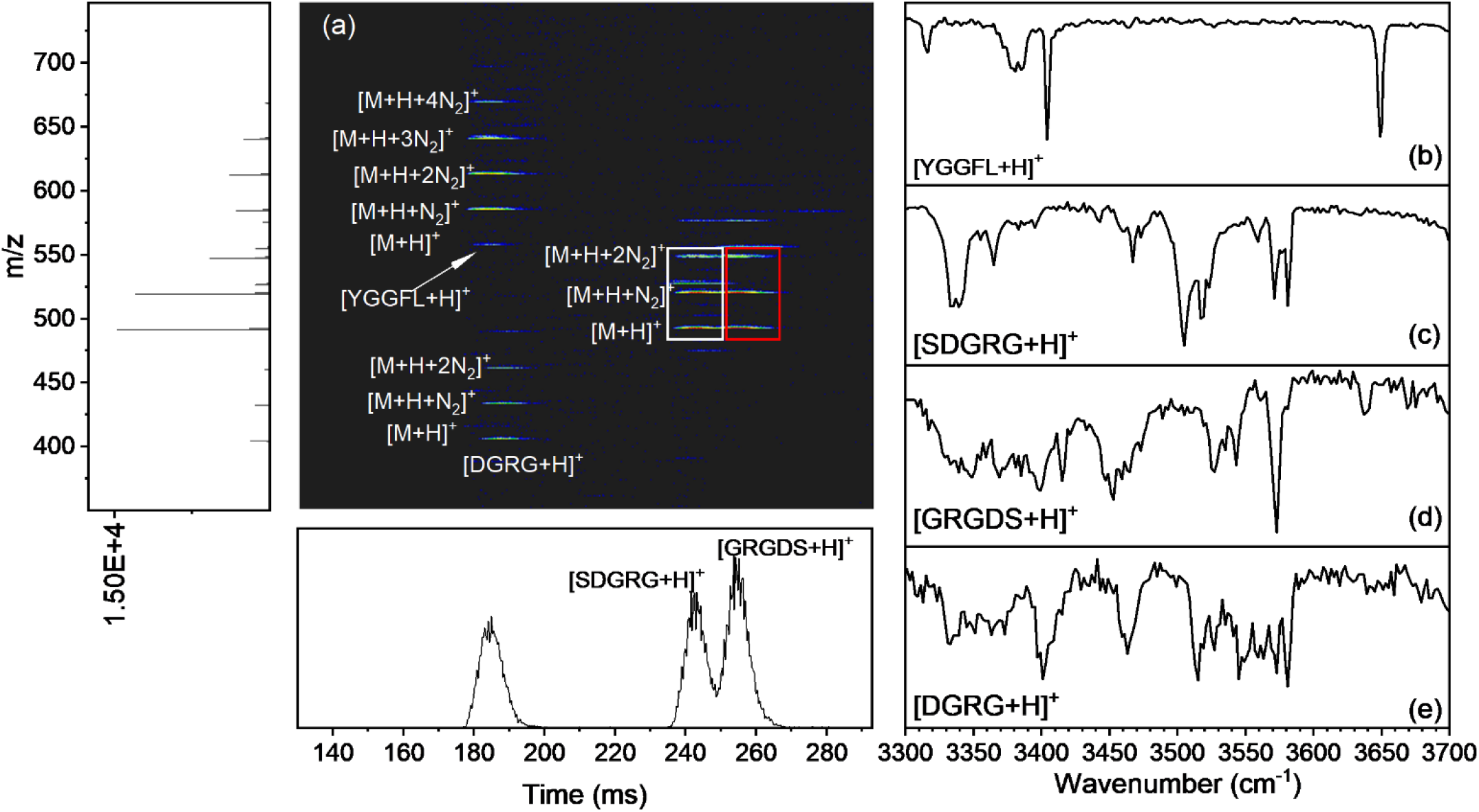
(a) Example of data collected while simultaneously recording IR spectra of all prominent ions in the sample. All peptide ions labeled appear mobility or mass resolved from one another and display 2 or more tags. The UIMF file is truncated to remove the 500 ms fill time. The resulting IR spectra of (b) [YGGFL+H]^+^, (c) [SDGRG+H]^+^, (d) [GRGDS+H]^+^, and (e) [DGRG+H]^+^.

In IR-only mode, no mobility separations are employed, and the ions are streamed through the 13-m SLIM as well as the cryo-SLIM while the cryo-SLIM TW amplitude is set to 0 V_pp_. Under these conditions, the ions continuously pass through the instrument and become thermalized and tagged as they pass through the cryo-SLIM. The ability to record IR spectra under these conditions is demonstrated in Figure 4. Figure 4a and 4b shows a 34-second extracted ion chromatogram for m/z 612.2 and 556.2 which corresponds to [YGGFL+2N_2_+H]^+^ and [YGGFL+H]^+^, respectively. The precursor ion (*m*/*z* 556.2) forms up to 3 tagged adducts with the doubly tagged species (*m*/*z* 612.2) being the most abundant under the chosen N_2_/He gas conditions for this experiment. The intensity of the MS data in Figures 4a and 4b is normalized to the maximum intensity of the [M+2N_2_+H]^+^ species, showing the tagged species is about 10x the abundance of the precursor ion. Figure 4c plots the ratio of the intensity of the 3 tagged species to the sum of the tagged species and the precursor ion. Note the intensity of all tagged species is considered when plotting the IR spectrum even though only the most abundant tagged species is shown in Figure 4a. The tagging efficiency is relatively high such that each time the laser becomes resonant with a vibrational transition, the formation of the precursor mass (Figure 4b) can clearly be observed. The data plotted in Figures 4a-4c were recorded using an average scan rate of 0.011 cm^-1^/ms and took about 34 seconds to scan over a 400 cm^-1^ region. Figure 4d shows the spectrum (red) recorded when the average scan rate is increased to 0.192 cm^-1^/ms, which took only ∼2 seconds to scan over the 400 cm^-1^ spectral range. For reference, the spectrum recorded at an average scan rate of 0.011 cm^-1^/ms is plotted in black for comparison. The FWHM of the transition near 3650 cm^-1^ is 4.0 cm^-1^ at a scan speed of 0.011 cm^-1^/ms and 8.1 cm^-1^ at a scan speed of 0.192 cm^-1^/ms indicating the loss in resolution at the higher scan speed. The loss in resolution appears marginal for this particular spectrum as there are only 5 transitions present in this spectral range. The most obvious difference between the two spectra is the two partially resolved bands between 3400 and 3350 cm^-1^ at a scan rate of 0.011 cm^-1^/ms (Figure 4d, black trace) appear as a single broader band when the scan rate is increased to 0.192 cm^-1^/ms (Figure 4d, red trace). While this is a small effect, care should be taken when the spectra become more congested. The IR spectra of [SDGRG+H]^+^ and [GRGDS+H]^+^ taken in IR-only mode at a scan rate of 0.011 cm^-1^/ms conditions are plotted in Figure 5a and 5b, respectively. The corresponding IR spectra recorded in IMS+IR mode taken at a scan rate of 1.0 cm^-1^/acquisition cycle are overlayed in grey for comparison. The IR spectra of both peptide ions taken in IR-only mode appear similar to those taken in IMS+IR mode showing the utility of this mode of operation to record IR spectra for mass-resolved ions quickly. When operating in IMS+IR mode, the length of a single acquisition cycle is limited by the time required for accumulation in addition to the IMS separation. Specifically, a single IR spectrum for the reverse peptide ions required a total time of ∼ 5 minutes to collect. In IR-only mode, this time is reduced to ∼ 34 seconds, as the limiting factor is now the laser scan rate. The ability to switch between the two operation modes is useful as a simple IMS experiment can be taken to evaluate whether isomeric species exist for a particular ion followed by the collection of IR spectra in IR-only mode if a single isomer is present or IMS+IR mode if multiple isomers are present. Further, the IR-only mode can be of utility if isomeric species are separated by LC prior to being ionized. In this scenario, the ability to quickly record IR spectra makes this technique more amenable to being coupled to LC.

**Figure 4.**
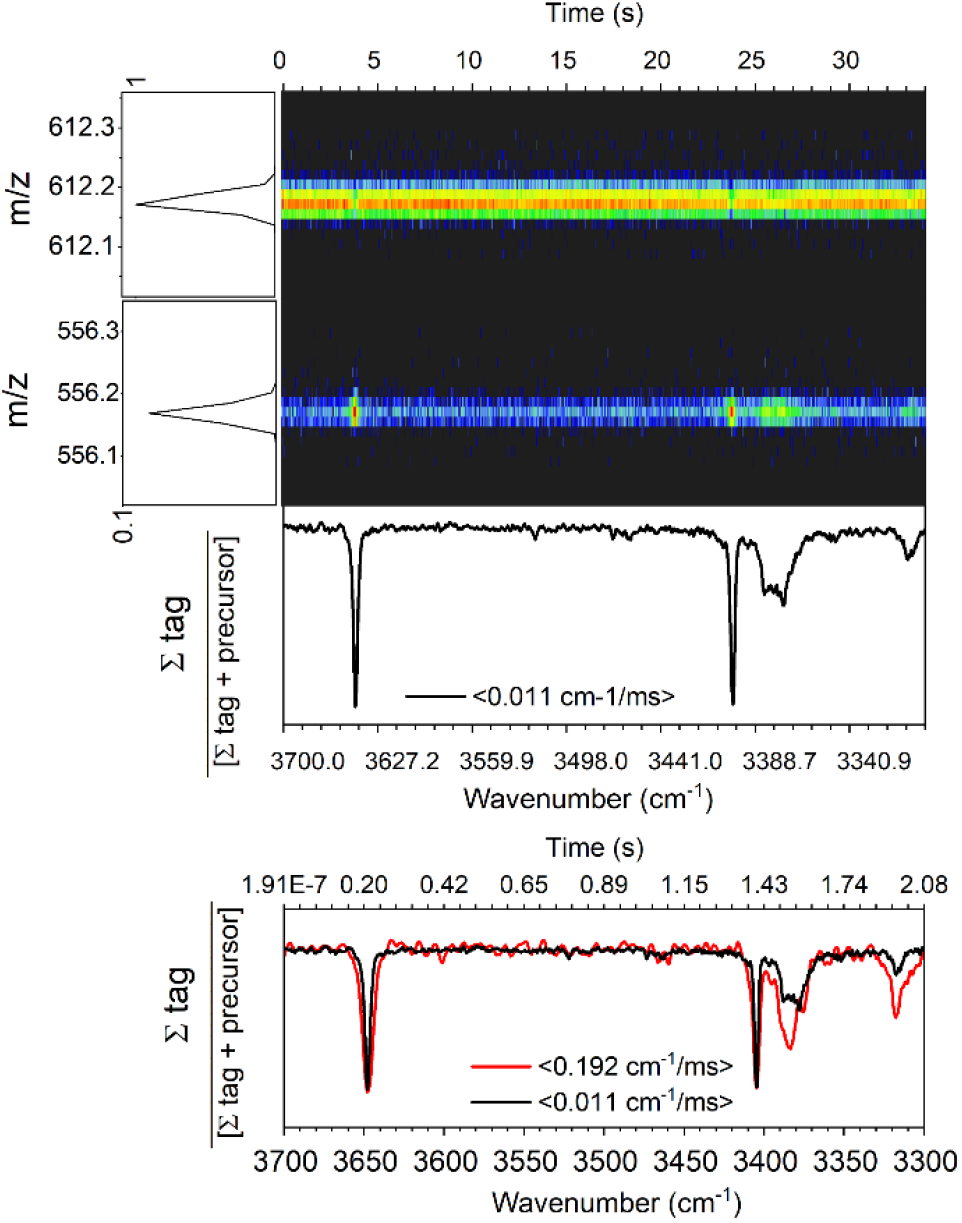
Extracted ion chromatograms for [YGGFL+H+2N_2_]^+^ (a) and [YGGFL+H]^+^ (b) in IR-only mode over the 34 second acquisition time period. (c) Resulting IR spectrum of [YGGFL+H]^+^ obtained by incorporating intensity from all tags (m/z 584, 612, and 640) using average scan rate of 0.011 cm^-1^/ms. (d) Resulting IR spectrum at an average scan rate of 0.192 cm^-1^/ms for a total acquisition time of ∼2 seconds (red), with the IR spectrum recorded at an average scan rate of 0.011 cm^-1^/ms overlayed for reference.

**Figure 5.**
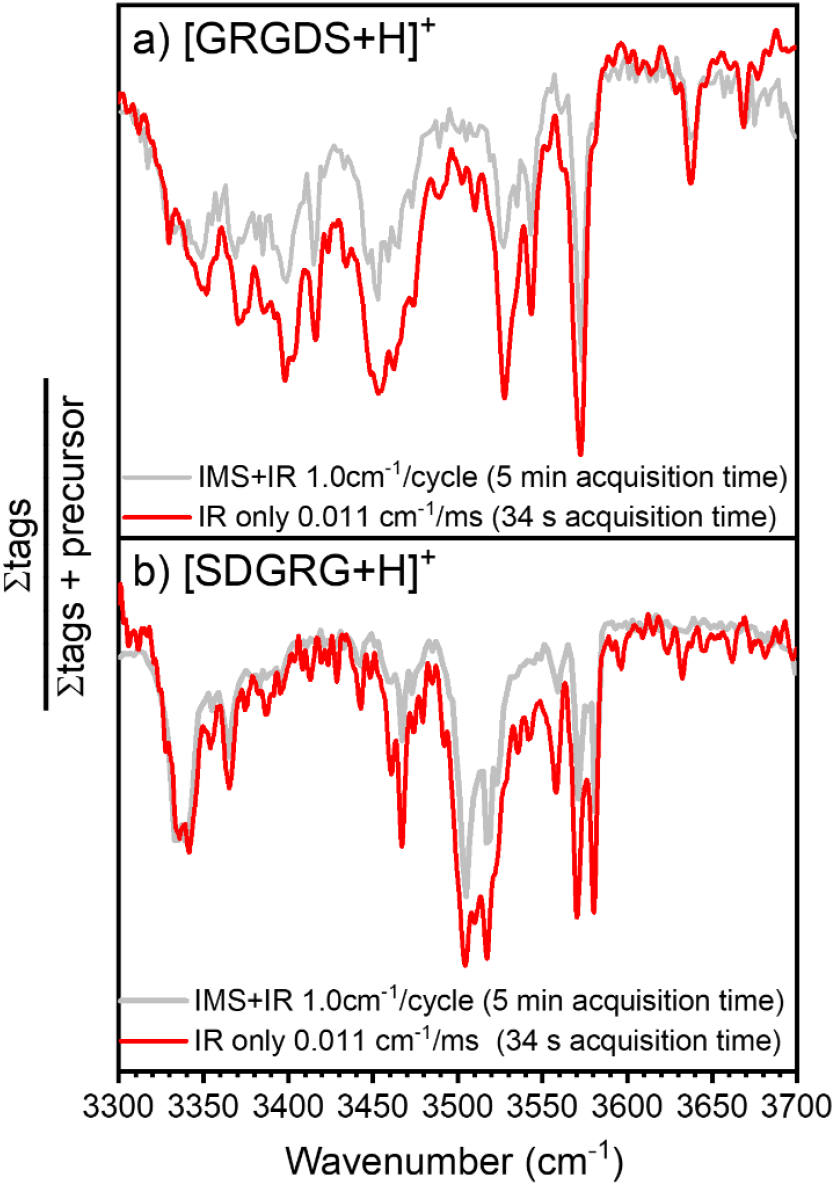
IR spectra of [GRGDS+H]^+^ (a) and [SDGRG+H]^+^ (b) recorded in IR only mode using an average scan rate of 0.011 cm^-1^/ms for a 34 s acquisition of their IR spectra (red traces). IR spectra recorded in IMS+IR mode are overlayed in grey for reference.

## Conclusion

We have developed an analytical platform adapted from an Agilent 6538 QTOF mass spectrometer. This platform combines high-resolution ion mobility spectrometry (IMS) using TWIMS-SLIM with a cryogenic messenger tagging technique. We demonstrated its capabilities using a small number of peptide ions. The key advancement of this platform is the ability to perform messenger tagging without trapping ions in the cryogenic SLIM trap. This allows for all mass or mobility-separated ions to be probed simultaneously in an efficient manner. Additionally, the platform enables the collection of IR spectra across a 400 cm^-1^ region in as little as 2 seconds when not performing IMS separations, showcasing its utility for high throughput sample characterizations. Calibrated CCS values were found to match reported values within 1%, and the IR spectrum of the benchmark YGGFL ion closely resembled previously published work. The precise description of 3-D shape through CCS and IR shows promise for distinguishing isomers and aiding in structural elucidation. Our future work will focus on leveraging these measurements for high-throughput characterizations and evaluating their potential for making de novo structural assignments when coupled with tandem mass measurements.

## Supporting information

Associate Content

## ASSOCIATED CONTENT

CCS calibration procedure, printed circuit board layout of the 90° constant field SLIM board, printed circuit board layout of the cryogenic SLIM, cross-section of the cryogenic SLIM trap assembly, example of the time sequence for the IMS+IR acquisition, cryo-TW SLIM guard effect on the tagged species, benchmark of the IR spectra of [YGGFL+H]^+^ to a previously published spectrum and comparison of the IR spectra recorded when a single vs. multiple tags are formed, IR spectra as a function of the laser step size.

## AUTHOR INFORMATION

## Author Contributions

The manuscript was written with contributions from all authors. All authors have approved the final version of the manuscript.

## Notes

The authors declare no financial interest.

## ACKNOWLEDGMENT

This research was supported by the PNNL Laboratory Directed Research and Development Program and is a contribution of the m/q initiative. C.P.H would like to acknowledge Bryson C. Gibbons, and Cameron Giberson for modifications the PNNL preprocessor and instrument control software. This project was performed in the Environmental Molecular Sciences Laboratory, a DOE Office of Biological and Environmental Research sponsored national scientific user facility located on the PNNL campus. Battelle operates PNNL for the DOE under contract DE-AC05-76RLO01830.

