## Supplementary material for "Development of a Platform for High-Resolution Ion Mobility Separations Coupled with Messenger Tagging Infrared Spectroscopy for High-Precision Structural Characterizations": Associate Content

**Table of Contents**

CCS Calibration Procedures

Figure S1: The printed circuit board layout of the constant field 90˚SLIM.

Figure S2: Printed circuit board layout of the cryogenic SLIM.

Figure S3: A cross-section view of the cryogenic TW SLIM trap assembly.

Figure S4: Example of time sequence for the IMS+IR acquisition.

Figure S5: Cryo-TW SLIM guard effect on the tagged species.

Figure S6: Benchmark of IR spectra of $[YGGFL+H]^+$ to previously published spectrum, comparison of IR spectrum recorded using a single tag vs. multiple tags.

Figure S7: IR spectra as a function of laser step size.

CCS Calibration:

Agilent tune mix ions 322-1822 were used as calibrant ions. Their arrival times were first plotted against their reduced CCS (Ω')

$$\left[\Omega = \Omega' z\sqrt{\mu^{-1}}\right] \qquad \text{S1}$$

where Ω is the known collision cross section, *z* is the charge, and μ is the reduced mass of the ion and $N_2$. The plot of arrival time vs. Ω' was then fitted using a power function of the form

$$y = a + bx^c \qquad \text{S2}$$

where *y* is the reduced CCS, *a* is the intercept, *b* is the slope, *x* is the arrival time taken at the peak apex, and *c* is the scaling factor. The coefficients (a, b, and c) from this fit were then used to solve for the reduced cross sections of the peptide ions based on their arrival times using equation S2 and then were converted to CCS by equation S1.

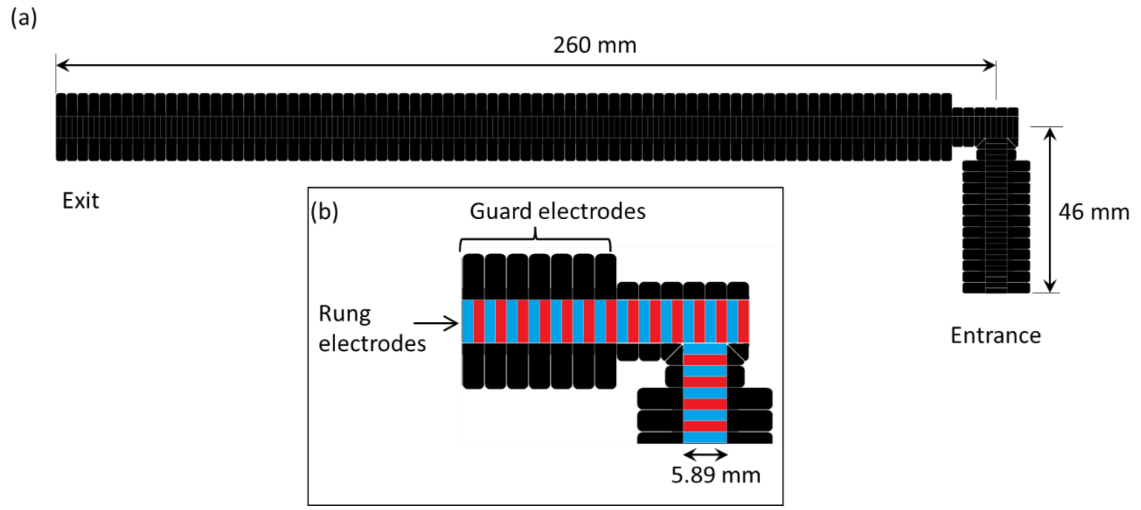


Figure S1: (a) The printed circuit board layout of the constant field 90˚SLIM showing dimensions. (b) A zoomed view of the 90˚ turn shows the guard electrodes (colored in black) and the rung electrodes (colored red and blue). A resistance chain connected to the rung electrodes provides a linear DC gradient between the entrance and exit electrodes. Adjacent rung electrodes have opposite phases of RF superimposed on the DC potentials, while guards only have DC. The guard electrodes are kept 5-10 V higher than the adjacent rung electrode to provide lateral confinement.

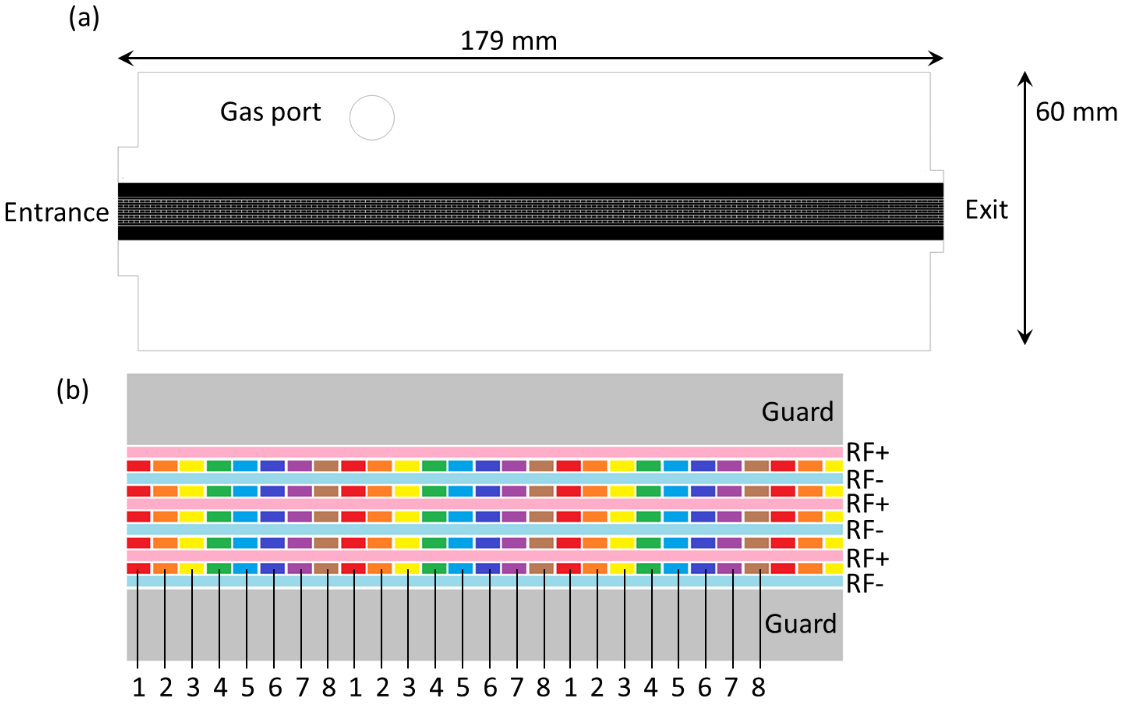


Figure S2. (a) Printed circuit board layout of the cryogenic SLIM showing the dimensions. (b) A zoom into a section of the SLIM tracks showing the TW, RF, and guard electrodes. A set of 8 segmented electrodes (numbered 1 to 8) are used to create the TW electric field.

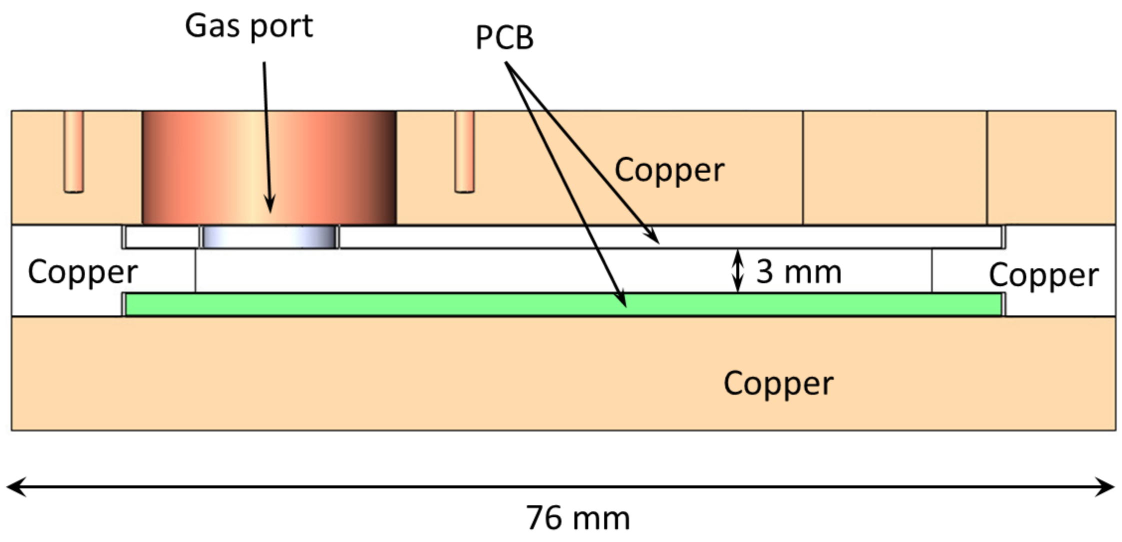


Figure S3. A cross-section view of the cryogenic TW SLIM trap assembly.

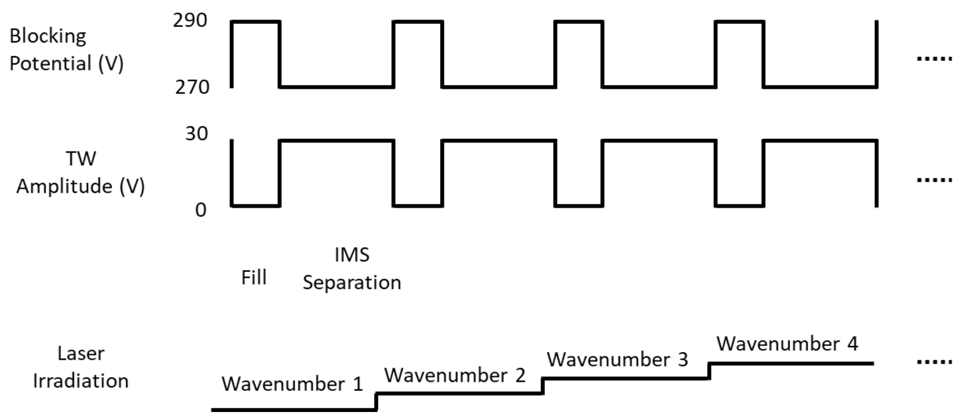


Figure S4: Example of time sequence for the IMS+IR acquisition.

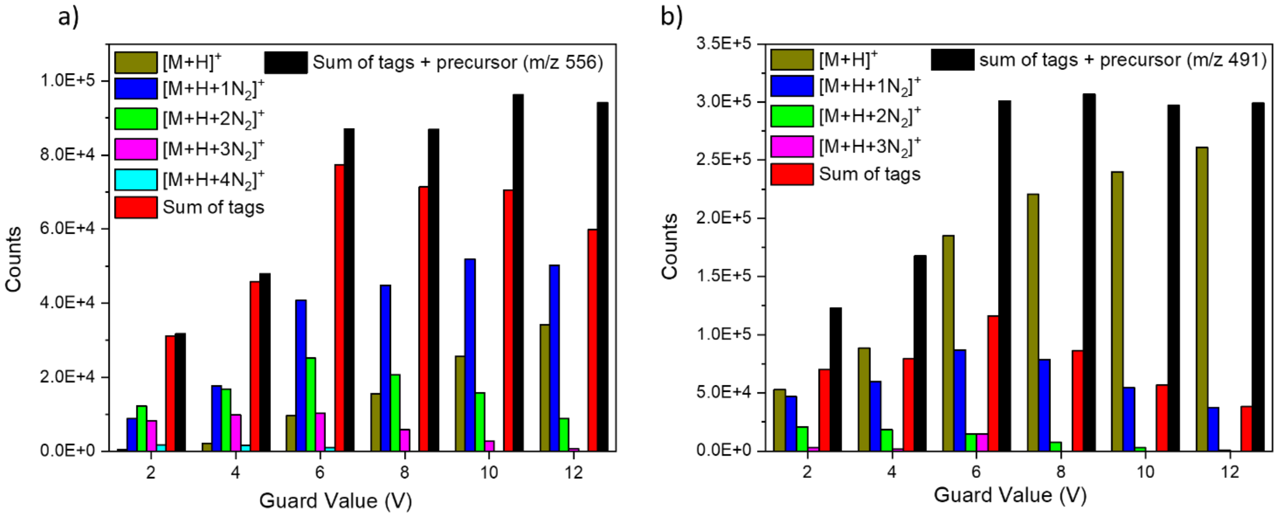


Figure S5: Ion intensities for precursor and associated tags for m/z 556 (a) and 491 (b) as a function of the guard voltage in the cryo-TW SLIM. Separation conditions in the 13m SLIM are consistent with that from Figure 3. The black bar (sum of tag and precursor) in each plot represents the total ion current that passes through the cryo-TW SLIM. This values plateaus around a guard voltage of 6 V. Below 6 V fewer ions are transmitted, and above 6 V, fewer tags are preserved.

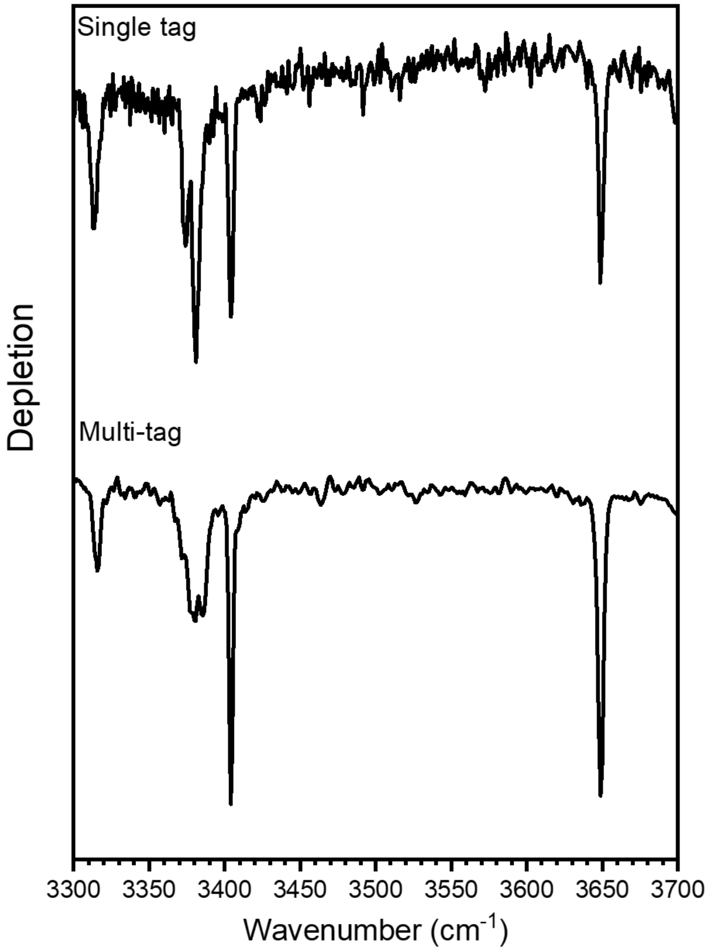


Figure S6: A reference IR spectrum for $[YGGFL+H]^+$ can be found in references 70 and 71 of the main text. The IR spectrum in the top panel is recorded when the $N_2$ gas conditions of the cryo-TW SLIM favor the formation of a single tagged species. This spectrum is highly similar to the spectra published in references 70 and 71. The IR spectrum in the bottom panel is recorded under typical conditions where multiple tags can be formed for $[YGGFL+H]^+$. The IR spectrum recorded when a single tagged and multiple tags are formed are also very similar, however, with a slight distortion of the two transitions between 3350-3400 $cm^{-1}$ indicating that the additional tags interact with these vibrational modes.

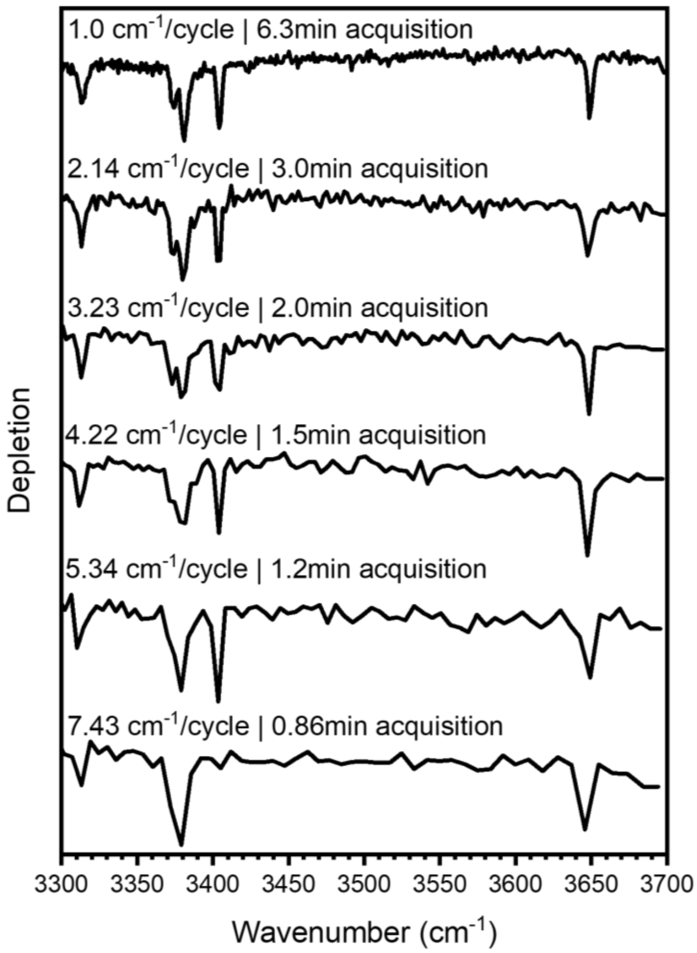


Figure S7: IR spectrum of $[YGGFL+H]^+$ as a function of the laser step size in IMS+IR mode. The separation TW and cryo-SLIM TW conditions are identical to those used in Fig. 3 of the main text but with an instrument cycle of 984 ms. The laser step size is increased from 1 $cm^{-1}$ to 7.43 $cm^{-1}$/acquisition cycle. A 3.23 $cm^{-1}$ / cycle step size appears to preserve the integrity of the IR spectrum while decrease the total acquisition time from 6.3 minutes to 2 minutes. Step size larger that ~3 $cm^{-1}$/cycle being to distort the IR spectrum as the resolution becomes worse.
